# Experiential diversity training and science learning for college students alongside peers with intellectual and developmental disabilities

**DOI:** 10.1101/2023.01.20.524901

**Authors:** Kaelin N. Rubenzer, Jonathan T. Pierce

**Affiliations:** Department of Neuroscience, The Center for Learning and Memory, Waggoner Center for Alcohol and Addiction Research, Institute of Neuroscience, University of Texas at Austin, Austin, TX, USA

## Abstract

Diversity, equity and inclusion (DEI) training can benefit STEM students. However, typical college settings often limit college students’ exposure to adult peers with intellectual and developmental disabilities (IDDs), a historically marginalized group. To lower this barrier, we developed a continuing education program, Lifelong Learning with Friends (LLWF), aimed at adults with IDDs on a large university campus, which provides diversity training to college students. In this program, undergraduate and graduate students from scientific and education disciplines are recruited to volunteer as peers and helpers. LLWF has reached hundreds of students with and without IDDs each year and more than 1,500 over the past 12 years. In our program, college students gain DEI training through learning sophisticated academic topics, including sciences, alongside adults with IDDs. Almost half (42%) of surveyed LLWF college volunteers did not have prior exposure to people with IDDs. Following program participation, we found that, irrespective of prior exposure, nearly all (98%) of volunteers had elevated their expectations of people with IDDs and reported increased interest in IDDs-focused research, education, social interaction, and advocacy. Additionally, college volunteers reported that they improved their science communication by seeing how science could be taught to a broad audience that includes adults with IDDs. We therefore suggest that other universities may consider our LLWF model to enhance DEI training by expanding opportunities for neurotypical students to befriend and learn science alongside adults with IDDs.

## Introduction

There is rising demand to expand the scope of training for future scientists, physicians, and science educators to incorporate real-world experiences through experiential diversity training [1]. Students beginning medical school often report feeling unprepared to relate to patients whose backgrounds or identity differ from their own [2–4]. This highlights the need to address STEM students’ inexperience outside of the classroom and in the real world. A major way to address this issue is by encouraging STEM students to seek experience with new and marginalized populations through paid or volunteer activities outside of the classroom. These experiences can be paradigm shifting and life changing; however, studies have shown that some current diversity training efforts need improvement. For one, they often are inconvenient because many are off campus and not for formal academic credit [5]. For those taught in the classroom, general diversity training aimed at STEM and medical students often relies on methods such as perspective taking, goal setting, or stereotype discrediting [6]. Simply discrediting socially unacceptable stereotypes can still inadvertently cause negative indirect biases [7–9]. Although these classroom methods can lay important groundwork for respecting and appreciating diversity, students likely benefit more from spending time with people from diverse backgrounds rather than reading about them [10,11].

Although both out of and in-classroom DEI training often focuses on critical diversity issues, such as race, gender, and socioeconomic status, most diversity equity and inclusion (DEI) training initiatives often overlook intellectual and developmental disabilities (IDDs). Many professionals entering medical and STEM fields report little to no prior formal training with people having IDDs or special education, despite people with IDDs representing a significant percentage of their future clientele [12–14]. This demographic is also particularly valuable for STEM students to interact with because of the growing percentage of college students and the professional workforce identified as neurodiverse. Growing awareness of broader neurodiversity proves the commonality of conditions like autism spectrum disorder (ASD), Down syndrome, and other IDDs, with an estimated 15-20% of the general population exhibiting some form of neurodivergence [15,16].

For decades, the Special Olympics has offered DEI experiences for the public to learn about people with IDDs. Special Olympics has two programs under their umbrella that have made strides in establishing mixer DEI experiences for college students: Unified sports programs and Best Buddies. Unified sports programs allow for athletes with and without disabilities to participate in organized athletic events and games together while also serving as a platform for building friendships, improving self-esteem, and promoting social acceptance among athletes without IDDs [17–21]. Similarly, Best Buddies is a nonprofit volunteer organization that promotes positive one-to-one interactions among high school and college students with and without IDDs, organizing monthly meetups, community hangouts, and routine communication via email, phone call, texting or social media [22,23]. Both Unified sport programs and Best Buddies utilize neurotypical participants to serve as similar age peers in athletic and social activities. However, recreationally inclusive programs may not succeed in demonstrating the level of what adults with IDDs are capable of learning and showing interest. This often-unappreciated goal is especially important to counter and better contextualize the “intellectual disability” label that assumes people with IDDs have no interest in or ability to learn sophisticated subjects in a college setting.

We sought to combine elements of diversity training with experiential learning with a novel platform, Lifelong Learning with Friends (LLWF), to connect mostly STEM and pre-medical college students to peers with IDDs. By offering academic material to college students, we offer a form of engagement with adults with IDDs that have unique strengths that complement recreational programs through Special Olympics. LLWF offers courses on a variety of diverse academic, cultural, and personal-development topics to adults with IDDs who learn alongside neurotypical college students in a nearly 2:1 reverse-inclusion format. Neurotypical college students volunteer in courses as class peers learning novel material while having the potential to gain meaningful experience interacting with people with IDDs. Typical DEI training is most often completed in short professional workshops, involving educational approaches like cultural humility training, or identifying bias [24]. LLWF is a unique way to achieve DEI training but through firsthand experience with people with IDDs rather than learning about them in a less personal workshop.

We wanted to determine if LLWF served as a valuable experiential diversity training opportunity for college student volunteers (hereafter referred to as volunteers). We surveyed volunteers before and after volunteering to assess their participation in LLWF as (1) an opportunity to meet and gain meaningful experience with people with IDDs, (2) a convenient and enjoyable volunteering experience, and (3) an inspiration for career development or advocacy in the disability field. We hypothesized that most volunteers would have little to no firsthand experience with peers with disabilities prior to volunteering due to limited opportunities for interaction. Because students who have little contact with peers with disabilities often overestimate the debilitating severity of disability [25,26] we also hypothesized that most volunteers would report little to no knowledge of adult peers with disabilities participating in age-appropriate activities, such as playing sports, taking courses, or having a job. Additionally, we hypothesized that most volunteers would rate their experience volunteering with LLWF as convenient, enjoyable, and transformative to raise their expectations of people with IDDs.

Finally, we wanted to determine if LLWF influenced some volunteers to pursue careers in fields related to IDDs.

### Lifelong Learning with Friends

Lifelong Learning with Friends (LLWF) is a reverse-inclusion college education program aimed at adults with IDDs that has been running continuously for 12 years at the University of Texas at Austin (UT). Reverse-inclusion classrooms recruit neurotypical students into special education settings to foster positive peer interaction, model age-appropriate behaviors, and offer academic support if needed [27,28]. LLWF utilizes the reverse-inclusion dynamic to allow for bi-directional social and academic learning between students with IDDs and volunteers. LLWF offers a range of courses over academic and recreational topics that are typical of a neurotypical college curriculum but are commonly unavailable to adults with IDDs. Adults with IDDs are able to select courses à la carte, allowing them to pick courses suited to their personal interests rather than adhering to more rigid degree plans typical of other PSE programs aimed at adults with IDDs. For comparison, the two other postsecondary education programs aimed at adults with IDDs in Austin (STEPS at Austin Community College and E4 Texas at UT Austin) offer more conventional training in remedial academics and independent life skills, paired with either general career training or personal attendant training, with limited options to customize their experience.

LLWF provides course topics that also appeal to volunteers, in part by introducing new material. Our science courses both complement and supplement their degree plan with topics that they otherwise would not have time to pursue. Volunteers are free to sign up for classes that appeal to their personal interests. Examples of past course topics include forensic science, pharmaceutical drug discovery, marine science, and more **(Figure 1)**. Courses are offered year-round in the evenings on weekdays and on Sunday afternoons, allowing college volunteers to easily fit them into their school schedules. Each course consists of six classes that meet once per week for 2-3 hours. Volunteers spend the entire class time with adults with IDDs.

**Fig 1.**
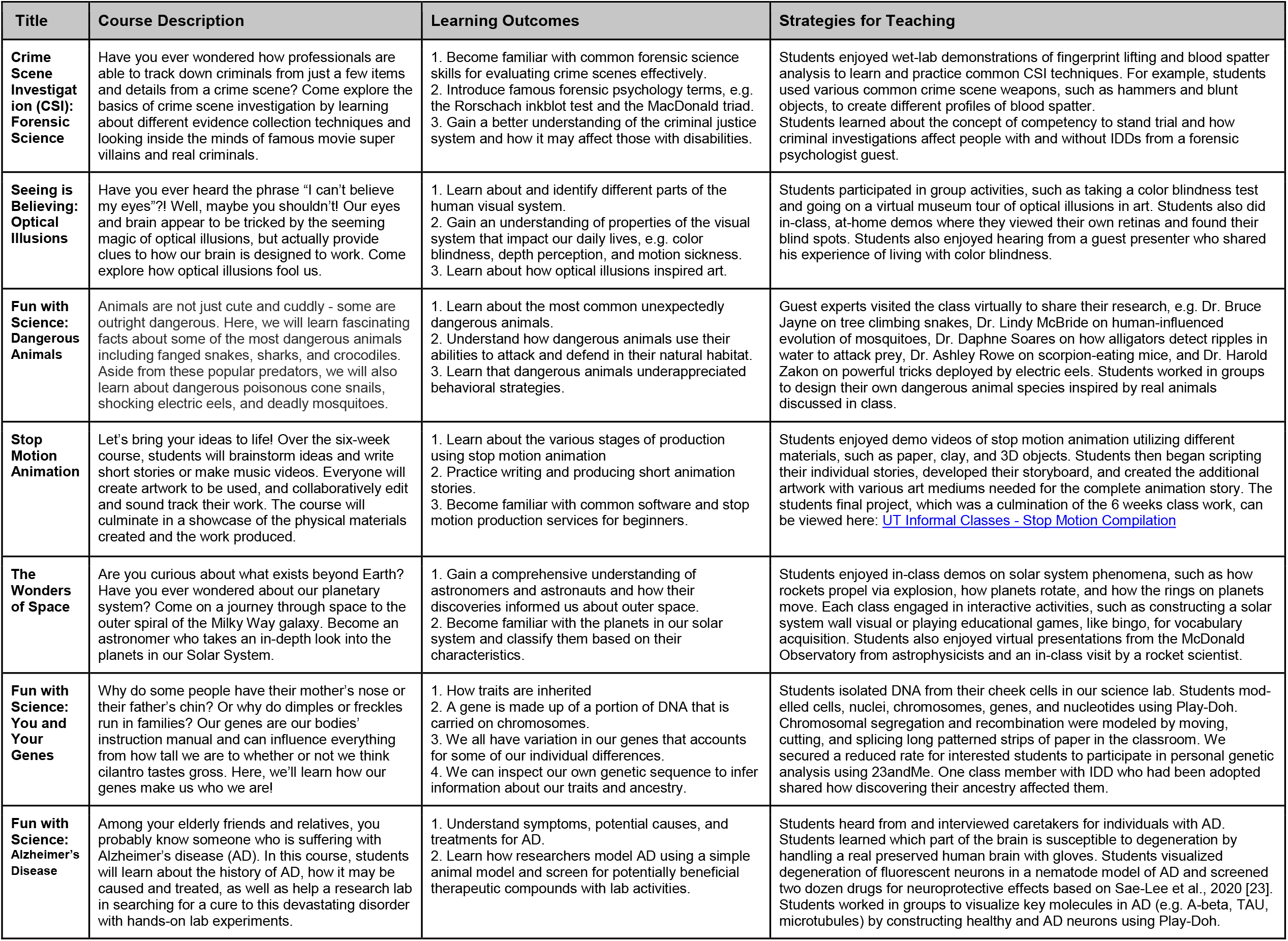
Previous LLWF science courses. LLWF courses include a range of sophisticated academic material typically covered at universities.

For some, it may be hard to imagine how reverse-inclusion courses with sophisticated topics including science can be effective for students with and without IDDs. To engage students with a wide range of abilities and different limitations, our instructors have employed learning strategies involving universal learning design (ULD) approaches. Some ULD methods we utilize in our science courses include *perceptible information* (using a variety of illustrations, precise simplified language, and written directions to present curriculum), *tolerance for error*, as well as *size and space for approach and use* (ensuring that size of projected or printed content is large enough and uncluttered) [29]. To keep adults with IDDs and volunteers engaged, our courses also feature hands-on interactive activities, wet-lab demonstrations featuring one-on-one support, independent inquiry, and expert guest lecturers. Examples of past LLWF courses are provided in **Figure 1**.

Public visibility of our students with IDDs on university grounds can demonstrate to volunteers and others that people with IDDs belong in higher education settings. All classes start at a campus cafe, where students with IDDs and volunteers converse freely for the first half hour. This social period presents a unique bidirectional learning exercise where adults with IDDs learn to reinforce their own age-appropriate behavior and social interactions with volunteers while volunteers learn first-hand about IDDs. Students then go to a nearby classroom in an adjacent building to engage interactively on the topic of the day. The classroom is on a floor with undergraduate science teaching labs, whereas upper floors house biomedical engineering, pharmacology, and toxicology research labs. Importantly, both the cafe and the classrooms are in an active and popular area, located in the heart of the campus, rather than in an area segregated from college students. By meeting at a popular cafe on campus and holding class in STEM buildings, our students with IDDs are better able to immerse themselves into university culture. Likewise, this on-campus location makes it convenient for STEM students to attend.

### Recruiting and training college volunteers

Undergraduate and graduate students, most of them interested in pre-med, neuroscience, and education fields, were recruited through university forums, list servs, and student organizations. Some students majoring in Education used LLWF participation as a required service-learning project (SLP) for their university course “Individual Differences”, which focuses on basic concepts, issues, and ways to accommodate people with disabilities. Other STEM students volunteered as an assignment in an upper level Neurogenetics course. Most volunteers, however, participate without course requirements. All volunteers are expected to attend every class, participate in classwork and discussions, and complete homework assignments.

During a 45-minute, mandatory orientation for all new volunteers, college students learn that they were expected to assume five roles during volunteering, regardless of course topic:

1. ***Mentor*** - When needed, college volunteers were encouraged by the instructor to assist students with IDDs in learning class topics. Examples include mediating small group activities, helping to scribe during worksheet assignments, amplifying or clarifying quieter or shyer students’ voices or written responses to enable participation in class discussion, or aiding in hands-on projects with students that required fine-motor skill support.
2. ***Peer*** - College volunteers are especially valuable in modeling age- and college-appropriate behavior, such as fostering reciprocal conversation, respecting personal space, basic classroom etiquette, and more. We have found that volunteers are in a unique position as peers to guide adults with IDDs towards age-appropriate behavior and learning that may be more effective than guidance from a parent or teacher figure because the adults with IDDs often want to emulate them [30,31].
3. ***Student*** - Volunteers have the unique opportunity to experientially learn about the science and social science of disability by befriending and learning alongside adults with IDDs. Many of our college volunteers are interested in the professional fields of medicine or education but may have limited opportunities to learn from people with IDDs, a potentially significant portion of their clientele base, outside of the classroom. Volunteering with LLWF allows college students to connect with and learn from adults with IDDs in real life instead of reading about them in class.
4. ***Friend*** - Many adults with IDDs are challenged at making friends due to difficulty in communication and/or relative isolation from similar-age peers. Like neurotypical people, social outlets outside of family and caretakers greatly enrich the quality of life for people with IDDs [10,32,33]. LLWF provides a valuable platform for adults with IDDs to re-enter the community and be introduced to college students similar to their age to form friendships.
5. ***Advocate*** - Volunteering with LLWF will hopefully change the volunteer’s expectations on the capabilities of people with IDDs. Volunteers are encouraged to bring what they learn about people with IDDs into their daily life at school and their future careers in medicine, education, and research.

Volunteers are trained to provide meaningful support for a variety of IDDs, such as one-on-one mentorship, redirecting to focus on class topics, or aiding in written assignments. Experienced course instructors (most often from special education fields) offer supplemental information regarding the social experience of volunteering, such as how to foster reciprocal conversation and support each student with unique IDDs. Instructors also lead a 10 minute debrief with volunteers after each class, which allowed volunteers to privately share their firsthand observations of IDDs. This debriefing period helps volunteers to better understand some of the behaviors typified by different IDDs, especially for volunteers inexperienced with people with IDDs. With the help of our experienced course instructors, volunteers were coached on how to provide disability-competent support. For instance, volunteers would have firsthand experience with repetitive behaviors or intense restricted interests typified by autism spectrum disorder, or speech intelligibility issues and impulse control common in some individuals with Down syndrome.

### Many college volunteers had little to no firsthand experience with people with IDDs

We sought to determine how many volunteers had firsthand experience with adults with IDDs prior to volunteering. We also sought to determine how many volunteers knew adults with IDDs that were involved with age-appropriate activities, like having a job, or playing sports, and if they had ever experienced people with IDDs being included in their classes. Over time, we administered pre-course surveys that evolved to include more questions, which accounts for the varied sample size (n = 112-131).

We hypothesized that many college students do not have experience with people with IDDs. Consistent with this idea, we found that we recruited two categories of volunteers. Nearly 54% of our volunteers reported having firsthand experience with people with IDDs (with 42% reporting having little to none) and 49% had previously socialized with people with IDDs (42% had not). **(Figure 2A and 2B)**.

**Fig 2.**
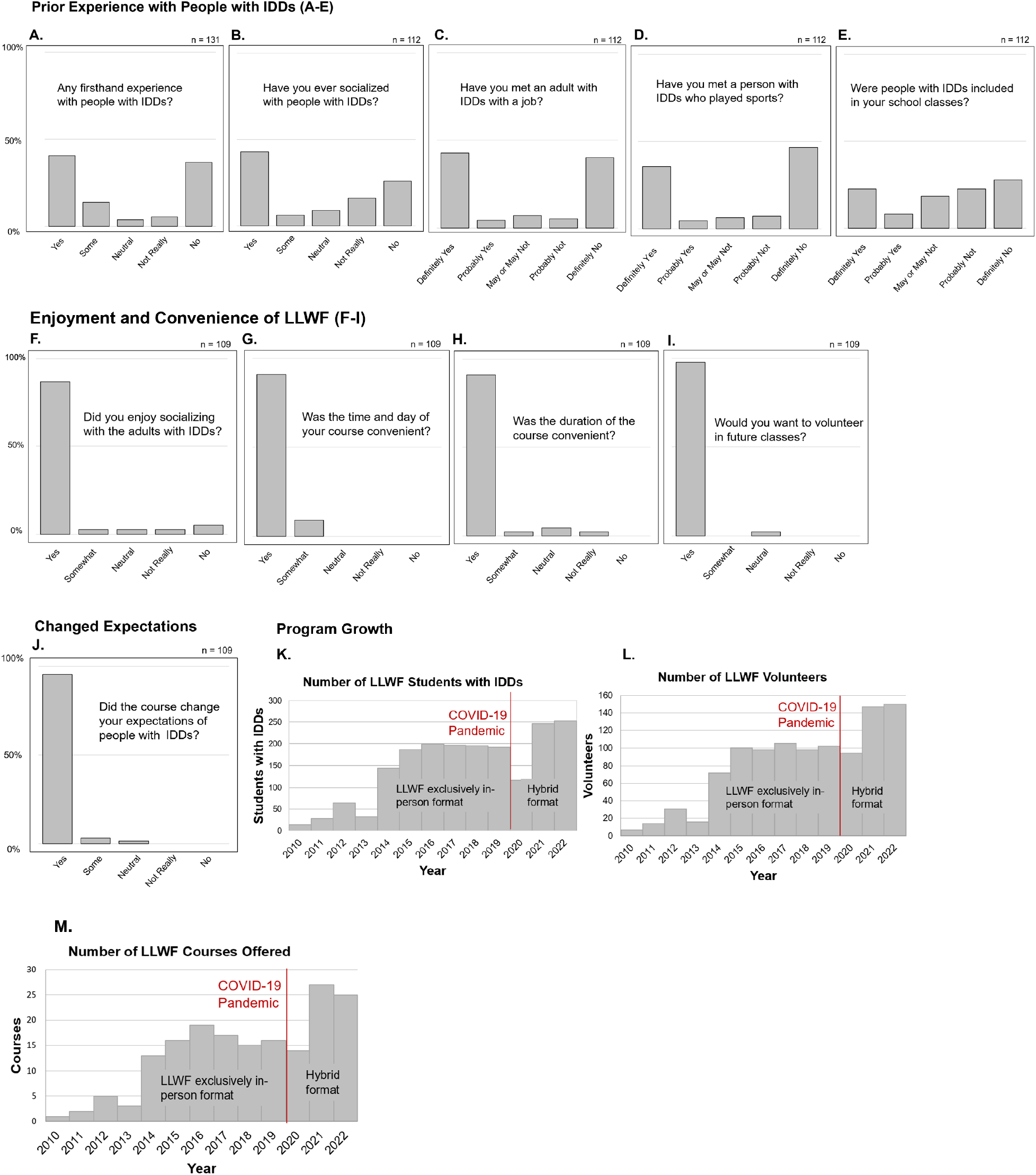
Pre- and post-course survey responses of volunteer participants and program growth over time. Prior to volunteering with LLWF, volunteers responded to several questions regarding their prior experience with people with IDDs **(A-E)**. In post-course surveys, volunteers were asked to rate enjoyment and convenience of LLWF volunteering **(F-J)**. In another portion of the post-course surveys, college students were asked if their expectations of people with IDDs changed after volunteering with LLWF **(I). \***LLWF has utilized slightly different versions of pre-class surveys as the program has expanded, which accounts for the variation in sample sizes across survey results. LLWF course and enrollment numbers since its inception in 2010 were analyzed to measure program growth over time **(K-M)**.

Many adults with IDDs participate in jobs and sports, but we hypothesized that most neurotypical people may not be aware of these activities and potential of adults with IDDs. Again, consistent with our hypothesis, we found that 47% of volunteers were aware that they had met an adult with IDDs with a job, 40% were aware that they knew a person with IDDs who played sports, and only 30% had experienced people with IDDs being included in their school classes (over 50% of our volunteers reported that they had likely not for each question) **(Figure 2C, 2D and 2E)**. Despite many volunteers having little experience with people with IDDs, we found that motivations for volunteering often involved a desire for helping and understanding people with IDDs **(Supplemental Figure 1)**.

### College volunteers reported LLWF as a convenient and enjoyable experience that bolstered their professional interests

In another portion of the post-course survey, volunteers were asked questions regarding enjoyment and the accessibility of volunteering with LLWF. We found that over 94% of surveyed volunteers enjoyed socializing with adults with IDDs, and almost all reported that the time, day, and duration of the course were convenient **(Figure 2F, 2G, and 2H)**. When asked if they wanted to volunteer in future courses, 98% responded “Definitely Yes” **(Figure 2I)**. When analyzing our volunteer signup system that dates to 2016, we found that we held a high volunteer retention rate, with 63% of volunteers returning for subsequent semesters (n = 470) even though there was no requirement to re-enroll. We also found that 32% of volunteers signed up for more than one course on different topics in the same semester. Because many of our volunteers are in the later stages of their undergraduate career, a natural phase out of the program is expected. Throughout our 12 years of operating LLWF, we have recruited and retained especially dedicated and enthusiastic volunteers who have volunteered for multiple classes each semester, stayed within LLWF for several years, or have transitioned to instructors for LLWF. Affectionately termed as “super volunteers”, LLWF has greatly benefited from these consistent and passionate volunteers.

We were interested in determining if volunteering with LLWF had positively influenced or intensified volunteers’ interest in fields of study related to IDDs, such as medicine, education or research. We included an optional free-response section on our post-class survey that asked volunteers if LLWF had any impact on their professional aspirations. We found that many students further developed their interests in medicine, education, and research by incorporating diverse perspectives and their positive experience volunteering **(Supplemental Figure 3)**.

### LLWF raised college volunteers’ expectations of adults with IDDs

We wanted to determine if volunteering in LLWF courses had changed volunteers’ expectations of adults with IDDs. Despite the bimodal distribution in terms of prior experience shown in Figure 2, we found a unimodal response with nearly all (98%) of students responding that the course had changed their expectations of people with IDDs (n=109) **(Figure 2J)**. We hypothesized that volunteers with little experience with people with IDDs would predictably raise their expectations after meeting students in our program because they had likely not met many adults with IDDs participating in challenging and age-appropriate activities, like working at a job, playing in sports, and participating and learning in sophisticated topics in a college classroom. For the group of volunteers that had prior experience with people with IDD, we hypothesized that their expectations still changed possibly because LLWF encourages adults with IDDs to learn sophisticated academic topics that are typically aimed at neurotypical adults, which is unique compared to most adult special education programs. To test these two hypotheses, we asked follow-up questions regarding volunteers’ change in expectations, represented in the table below **(Supplemental Figure 2)**. Many discussed surprise about being able to relate with students with IDDs, or were impressed that adults with IDDs were able to participate meaningfully during class.

## DISCUSSION

An important outcome of LLWF is the benefits the STEM volunteers receive. The opportunity to meet and befriend adult peers with IDDs may benefit their careers by galvanizing IDDs-focused research and care or inspiring them to actively seek experience with neurodiverse groups. Common STEM volunteer opportunities, such as shadowing a physician or working within marginalized communities, can be passive learning, time-intensive, require off-campus transportation, and interfere with the demanding class schedule of college students. Competitive pre-medical and STEM students with career-related motivations for resume-building can capitalize on LLWF serving as a convenient and interactive experiential diversity training experience located in the heart of campus. UT students often use LLWF as a mandatory sanctioned voluntary activity for their for-credit classes, such as the education course “Individual Differences” and an upper-level neuroscience lab course about neurogenetics. Additionally, volunteering in programs such as LLWF satisfies the growing interest in implementing diversity, equity, and inclusion principles to diversify medical and STEM fields [34]. DEI training can be an important force for reducing health care disparity, improving patient outcomes, and reducing the communication barrier between scientists and physicians with the general public [35,36].

Volunteering in programs like LLWF may help establish a much-needed dialogue between STEM students and neurodivergent audiences. By fostering students’ interpersonal communication, it encourages a more balanced STEM education experience through incorporating diverse perspectives and liberal arts teaching methods [37–39]. By design, college STEM training often lacks integration of liberal arts and humanities, which may limit STEM students’ ability to adopt different pedagogies and delivery methods to relate to others outside of their field. It is the responsibility of aspiring physicians, researchers, and educators to have strong communication skills to foster the public understanding and appreciation of science [40]. Many assume that STEM experts in their field should naturally be able to communicate well to a layperson audience, but the increasing specialization, conventions of academic publishing and presentations, and lack of formal communication training result in the ever-growing gap between scientists and the public [41,42]. Effective communication in science requires training and ample practice. DEI programs like LLWF incorporate diversity perspectives by engaging college students with people with IDD, encouraging them to share university space and begin practicing accommodating communication styles, satisfying equity and inclusion ideals. Serving as class peers and providing one-on-one support to adults with IDDs allows volunteers to practice effective ways to communicate with an atypical audience.

The limited contact between students with and without disabilities throughout schooling has likely led to negative attitudes towards the inclusion of students with IDDs in academic settings. Studies have shown that young students with limited contact are more likely to 1) perceive students with IDDs as more impaired than reality, 2) believe that students with IDDs should only participate in nonacademic classes, and 3) avoid social interaction with a peer with IDDs, particularly outside of school [25,26]. Moreover, the “intellectual disability” label, while chosen to replace prior offensive terms, still conveys low ability and interest to learn sophisticated subjects [43]. Without meaningful exposure to people with IDDs, these negative stereotypes and low expectations of people with IDDs learned through most American school systems and societies have limited opportunity to be rectified [44,45]. An opportunity for encouraging exposure and inclusivity is missed by the exclusion of peers with IDDs in formal education settings. Despite the growing movement for inclusive learning [46], many people with IDDs remain segregated to special education classrooms, often with restrictive academic curriculum and limited exposure to neurotypical peers [47–50]. By design, postsecondary education options that are historically aimed at neurotypical students have been generally inaccessible to adults with IDDs. Although the development of organizations, including Think College, are connecting more adults with IDDs with opportunities for inclusive higher education, many adults with IDDs have limited options to be included in age-appropriate college settings without having to be largely independent, technology savvy, or to have consistent access to transportation [51].

Greater attention is needed regarding post-high school outcomes and quality of life for adults with IDDs. Many adults with IDDs have limited chances for social interaction, poor vocational options, lower rates of postsecondary education involvement, and lack of transportation [52,53]. Compounding factors, such as financial hardship, lack of consistent and quality IDDs-trained staffing, and poor community support likely contribute to the often unfortunate professional and social outcomes for adults with IDDs [54]. Improving social networks for people with IDDs has far reaching benefits. For instance, expanding social personal networks of people with Down syndrome can mitigate cognitive decline and the development of Alzheimer’s disease [55].

Many IDDs-transition specialists suggest that coordination with either postsecondary education institutions or local agencies that provide employment support can provide bridges to a more self-determined and secure future [52,53,56]. LLWF presents opportunities for adults with IDDs to make adult friends, pursue their interests in formal higher education settings, and encourage inclusion and advocacy among neurotypical peers. In the future, we hope to determine to what extent LLWF has directly improved post-high school outcomes and quality of life for adults with IDDs in a longitudinal study.

### Factors to initiate, sustain and replicate LLWF

The successful initiation and development of LLWF over a decade ago relied on six key factors:

1. A founder or director with the passion and drive to develop a program that provides a college-level education to adults with IDDs
2. A sponsor for the program to take place
3. Instructors, preferably with experience in special education or IDDs services
4. College students interested in volunteering as class peers
5. Adults with IDDs and their caretakers interested in postsecondary education options

We found that there are benefits of having a tenure-track professor initiate LLWF on our university campus. A tenured professor may have access to reserve classrooms on campus, such as conference rooms and science labs, that an outside instructor or non-tenure track professor may not. Professors also may more easily recruit peer professors, graduate students, and postdocs to serve as guest lecturers, which help fortify learning in our courses. In turn, college professors may use initiation of a similar program to LLWF as an impactful outreach project in education and science grants (e.g. National Science Foundation, Beckman Foundation, and Howard Hughes Medical Institute). If potential program founders have family members with IDDs, then they may also gain additional assistance from their support organizations by providing a more compelling personal account to develop the program when communicating to people unfamiliar with IDDs.

Replicating our program may be particularly successful if done in collaboration with the school’s neuroscience department, as there is rapid expansion of neuroscience undergraduate programs. As a tenure-track professor in the neuroscience department, our program founder was well-positioned to recruit STEM undergraduate students interested in volunteer opportunities. Because conditions like ASD, Down syndrome, Williams syndrome and many other IDDs directly impact neurodevelopment and function, neuroscience students interested in cognitive development and function have the unique opportunity to experientially learn about these disorders directly from adults with IDDs. We attribute much of our success in recruiting and retaining volunteers due to the rapid growth of majors relevant to IDDs study and service. From 2017 to 2022, neuroscience represented one of the fastest growing majors at UT with a growth rate of 50% (about 750 to over 1100 students). Additionally, outreach to similar IDDs support organizations in Texas, such as Down Syndrome Association of Central Texas (DSACT), Autism Society of Texas, Adults Independent and Motivated (AIM), and Best Buddies, have helped advertise LLWF to potential students in and outside of Austin.

### Continuation and adaptation of LLWF during Covid

In response to the global COVID-19 pandemic, LLWF pivoted to offer virtual courses for the first time beginning in the spring semester of 2020. Since its inception in 2010, LLWF exclusively operated as an in-person program hosted on UT campus. Due to CDC protective guidelines and university closure, LLWF discontinued in-person courses in consideration of adults with IDDs in spring 2020. Adults with IDDs are considered a high-risk group for COVID-19 secondary complications, exhibiting 11 times higher odds of dying from COVID-19 in the first two months of the pandemic compared to non-disabled people [57,57]. For example, people with DS in general often exhibit a higher prevalence of respiratory tract infections and immune dysregulation, which makes them especially susceptible to serious complications and potential mortality from COVID-19 [58–60]. Additionally, many adults with IDDs live within residential group homes or with elderly family members, which presents the additional threat of community-based spread [61–64]. To mitigate the spread of infection, many adults with IDDs became homebound, with limited opportunities to socialize and participate in the community. Unlike many neurotypical people their age, people with IDDs are often less technology-savvy and participate less on social media platforms and online based communities, further limiting their means of socialization beyond live-in family members for the duration of the pandemic [59,65– 70].

Although switching to an online format was done out of necessity, the transition proved to be a serendipitous opportunity for LLWF to expand. For the first time, students and volunteers far from Austin were able to join our program, with volunteers and students from over 11 states and 22 universities now regularly attending courses to date. Since the successful implementation of a hybrid alternate online or in-person format in late 2020, LLWF has offered a greater number of courses offered per year and reached more students with IDDs and LLWF volunteers than ever before **(Figure 2K, 2L and 2M)**. From interactions with new adults with IDDs and their families, we also found that an option for virtual PSE is especially attractive to those who are wheelchair bound, have limited access to transportation, or live in smaller areas without many IDDs services. For instance, students who are bedbound due to muscular dystrophy have participated in our online courses using eye-directed communication software. Additionally, the online format encouraged adults with IDDs to become more technology-savvy, with many adults with IDDs now comfortable navigating Zoom online classrooms, performing independent internet searches, creating basic PowerPoint presentations, and utilizing online resources like YouTube for educational purposes. Given that Covid and related illnesses will continue to circulate and present greater risk to certain adults with IDDs and their families, bolstering in-person courses with an online option may represent a working strategy to replicate programs like LLWF at additional colleges.

### Limitations

Although an important mission of LLWF is to provide volunteers the opportunity to witness adults with IDDs be afforded the dignity to take courses on mature topics like their college peers, we found that many families were reluctant to enroll their students with IDDs in certain course topics that they reported seemed controversial, complex, or too mature. Courses that were popular with neurotypical college students, such as “Psychology of Science Fiction”, “Fun with Science: Viruses”, “Fun with Cultures: World Religions”, and “Art of Frida Kahlo” often received below the minimum enrollment requirement from adults with IDDs (6 students). Nevertheless, we succeeded in securing better enrollment on some courses with mature topics by later re-marketing them with more lighthearted titles and descriptions in later semesters. For instance, a course originally focused on business and marketing gained better enrollment in subsequent semesters when rebranded as the more palatable “History of Walt Disney”, and a course on intimacy and consent was retitled as “Romance in the Movies”. We also merged previously unsuccessfully drier topics, such as math or etiquette, under course topics with a more fun context. For example, whereas math and etiquette were unpopular courses, we found that a “It’s My Party!” course that combined these topics was popular since students worked together to budget and plan a catered banquet in the UT Tower.

When LLWF originated on UT campus over a decade ago, we struggled at times to achieve societal acceptance and buy-in from the university to host an inclusive education program on campus. Some administrators and university services were reluctant to support LLWF because several university policies inadvertently created barriers to the establishment of LLWF on campus. For instance, policy dictates that many campus rooms and equipment are reserved for exclusive use by undergraduates in certain majors, but not to the public outside of the university. While basic policies like this are important to the general protection of university spaces, they inadvertently restrict adults with IDDs from integrating into typical college settings unless they have strong ties to an in-group advocate. To gain university buy-in, we reframed our requests, explaining that our program primarily benefits university students by teaching them about disabilities. Conversely, certain organizations off campus, such as restaurants and some guest lecturers were not interested in our program if we advertised ourselves as mostly helping college students. To surmount this problem, emphasizing that our program benefits students with IDDs spurred their involvement. These problems and solutions demonstrate how LLWF’s reverse-inclusion format can offer an unexpected strategy to gain wider buy-in. Better understanding of the impact of LLWF can be achieved through further evaluations. Because the authors are involved with LLWF, future studies by outsider participants with either expertise in continuing education or scientific outreach programs would be valuable.

## CONCLUSION

LLWF presents a new way to add to inclusive education options on campus, while providing a valuable experiential volunteering activity that instills core principles of DEI training in college students. Providing more opportunities for inclusive higher education not only improves social and professional outcomes of adults with IDDs, but also can encourage a more inclusive society by improving the expectations and impressions of people with IDDs. Inclusive education (IE) programs on neurotypical college campuses, like LLWF, benefit both neurotypical and students with IDDs alike. IE programs encourage neurotypical students to learn alongside students with IDDs, improve their academic communication skills, and enhance societal acceptance of people with IDDs. IE programs like LLWF may be readily implemented at other higher education institutions to enhance inclusivity in academic settings and provide a valuable platform for neurotypical students to befriend and learn alongside adults with IDDs.

## ACKNOWLEDGEMENTS

We appreciate thoughtful discussions and feedback on this manuscript by Joshua Russell, Sophie Sanchez, and Haiti Qin. Statistics on undergraduates majoring in Neuroscience at UT were helpfully compiled by Kathryn Hendren.

## REFERENCES

1. Szczurek K, Furgał N, Szczepanek D, Zaman R, Krysta K, Krzystanek M. “Medical Student Syndrome”—A Myth or a Real Disease Entity? Cross-Sectional Study of Medical Students of the Medical University of Silesia in Katowice, Poland. IJERPH. 2021;18: 9884. doi:10.3390/ijerph18189884

2. Faulconer EK, Wood B, Griffith JC. Infusing Humanities in STEM Education: Student Opinions of Disciplinary Connections in an Introductory Chemistry Course. J Sci Educ Technol. 2020;29: 340–345. doi:10.1007/s10956-020-09819-7

3. Davis DLF, Tran-Taylor D, Imbert E, Wong JO, Chou CL. Start the Way You Want to Finish: An Intensive Diversity, Equity, Inclusion Orientation Curriculum in Undergraduate Medical Education. Journal of Medical Education and Curricular Development. 2021;8: 238212052110003. doi:10.1177/23821205211000352

4. Chapman EN, Kaatz A, Carnes M. Physicians and Implicit Bias: How Doctors May Unwittingly Perpetuate Health Care Disparities. J GEN INTERN MED. 2013;28: 1504–1510. doi:10.1007/s11606-013-2441-1

5. Cormack I, Konidari S. Integrating Volunteering with the Curriculum: present initiatives and future possibilities. London Met Repository. 2007;4: 89–97.

6. Lindsey A, King E, Hebl M, Levine N. The Impact of Method, Motivation, and Empathy on Diversity Training Effectiveness. J Bus Psychol. 2015;30: 605–617. doi:10.1007/s10869-014-9384-3

7. Dobbin F, Kalev A. Why Doesn’t Diversity Training Work? The Challenge for Industry and Academia. Anthropology Now. 2018;10: 48–55. doi:10.1080/19428200.2018.1493182

8. Corrigan PW, Penn DL. Lessons from social psychology on discrediting psychiatric stigma. American Psychologist. 1999;54: 765–776. doi:10.1037/0003-066X.54.9.765

9. Berinsky AJ, Mendelberg T. The Indirect Effects of Discredited Stereotypes in Judgments of Jewish Leaders. Am J Political Science. 2005;49: 845–864. doi:10.1111/j.1540-5907.2005.00159.x

10. Thomas A, Bax M, Smyth D. The health and social needs of young adults with physical disabilities. Cambridge University Press; 1989.

11. Clark-Ibanez M, Felmlee D. Interethnic Relationships: The Role of Social Network Diversity. J Marriage and Family. 2004;66: 293–305. doi:10.1111/j.1741-3737.2004.00021.x

12. Cooc N, Yang M. Diversity and Equity in the Distribution of Teachers With Special Education Credentials: Trends From California. AERA Open. 2016;2: 233285841667937. doi:10.1177/2332858416679374

13. Theoret C, Patel R, Thangamathesvaran L, Shah R, Chen S, Traba C. Creating Disability-Competent Medical Students Via Community Outreach. Journal of the National Medical Association. 2021;113: 69–73. doi:10.1016/j.jnma.2020.07.010

14. Ryan TA, Scior K. Medical Students’ Attitudes Towards Health Care for People with Intellectual Disabilities: A Qualitative Study. J Appl Res Intellect Disabil. 2016;29: 508–518. doi:10.1111/jar.12206

15. Doyle N. Neurodiversity at work: a biopsychosocial model and the impact on working adults. British Medical Bulletin. 2020;135: 108–125. doi:10.1093/bmb/ldaa021

16. Maenner MJ, Shaw KA, Bakian AV, Bilder DA, Durkin MS, Esler A, et al. Prevalence and Characteristics of Autism Spectrum Disorder Among Children Aged 8 Years — Autism and Developmental Disabilities Monitoring Network, 11 Sites, United States, 2018. MMWR Surveill Summ. 2021;70: 1–16. doi:10.15585/mmwr.ss7011a1

17. Sherrill C. Adapted physical activity, recreation, and sport: crossdisciplinary and lifespan. 5th ed. Boston, Mass: WCB/McGraw-Hill; 1998.

18. Vickerman P. Teaching physical education to children with special educational needs. New York, NY: Routledge; 2007.

19. Teodorescu S, Bota A, Stănescu M. Activităţi fizice adaptate-pentru persoanele cu deficienţe senzoriale, mintale şi defavorizate social. Ed Printech. Bucureşti: 4–15.

20. Siperstein GN, Harada CM, Parker RC. Comprehensive National Study of Special Olympics Programs in the United States. University of Massachusetts Boston. 2005;A Special Report.

21. Bota A, Teodorescu S, şerbănoiu S. Unified Sports – A Social Inclusion Factor in School Communities for Young People with Intellectual Disabilities. Procedia - Social and Behavioral Sciences. 2014;117: 21–26. doi:10.1016/j.sbspro.2014.02.172

22. Hardman ML, Clark C. Promoting Friendship Through Best Buddies: A National Survey of College Program Participants. Kliewer C, editor. Mental Retardation. 2006;44: 56–63. doi:10.1352/0047-6765(2006)44[56:PFTBBA]2.0.CO;2

23. Lehman BJ. Evaluating the “Best Buddies” program: The influence of friendships on attitudes towards students with developmental disabilities. The Claremont Graduate University. 2002.

24. Corsino L, Fuller AT. Educating for diversity, equity, and inclusion: A review of commonly used educational approaches. J Clin Trans Sci. 2021;5: e169. doi:10.1017/cts.2021.834

25. Siperstein GN, Parker RC, Bardon JN, Widaman KF. A National Study of Youth Attitudes toward the Inclusion of Students with Intellectual Disabilities. Exceptional Children. 2007;73: 435–455. doi:10.1177/001440290707300403

26. Bunch * G, Valeo A. Student attitudes toward peers with disabilities in inclusive and special education schools. Disability & Society. 2004;19: 61–76. doi:10.1080/0968759032000155640

27. Elaldi ş, ÇiFçi T, YerliYurt NS. Tersine Kaynaştirmaya Genel Bakiş: Nitel Bir Çalişma. OPUS Uluslararasi Toplum Araştirmalari Dergisi. 2021;17: 788–812. doi:10.26466/opus.818118

28. Schoger KD. Reverse Inclusion: Providing Peer Social Interaction Opportunities to Students Placed in Self-Contained Special Education Classrooms. Teaching Exceptional Children-Plus 2. 2006.

29. King-Sears M. Universal Design for Learning: Technology and Pedagogy. Learning Disability Quarterly. 2009;32: 199–201. doi:10.2307/27740372

30. Winstead O, Lane JD, Spriggs AD, Allday RA. Providing Small Group Instruction to Children With Disabilities and Same-Age Peers. Journal of Early Intervention. 2019;41: 202– 219. doi:10.1177/1053815119832985

31. McWilliam RA, Bailey DB. Effects of Classroom Social Structure and Disability on Engagement. Topics in Early Childhood Special Education. 1995;15: 123–147. doi:10.1177/027112149501500201

32. Francis GL, Stride A, Reed S. Transition strategies and recommendations: perspectives of parents of young adults with disabilities: Transition strategies. British Journal of Special Education. 2018;45: 277–301. doi:10.1111/1467-8578.12232

33. Raymond É, Grenier A, Hanley J. Community Participation of Older Adults with Disabilities: Community participation and disabilities. J Community Appl Soc Psychol. 2014;24: 50–62. doi:10.1002/casp.2173

34. Rosenkranz KM, Arora TK, Termuhlen PM, Stain SC, Misra S, Dent D, et al. Diversity, Equity and Inclusion in Medicine: Why It Matters and How do We Achieve It? Journal of Surgical Education. 2021;78: 1058–1065. doi:10.1016/j.jsurg.2020.11.013

35. Campbell FK. Medical Education and Disability Studies. J Med Humanit. 2009;30: 221– 235. doi:10.1007/s10912-009-9088-2

36. Simms C. Voice: the importance of diversity in healthcare. Int J Clin Pract. 2013;67: 394–396. doi:10.1111/ijcp.12134

37. Committee on Integrating Higher Education in the Arts, Humanities, Sciences, Engineering, and Medicine, Board on Higher Education and Workforce, Policy and Global Affairs, National Academies of Sciences, Engineering, and Medicine. The Integration of the Humanities and Arts with Sciences, Engineering, and Medicine in Higher Education: Branches from the Same Tree. Skorton D, Bear A, editors. Washington, D.C.: National Academies Press; 2018. p. 24988. doi:10.17226/24988

38. Gorbaneva V, Shramko L. Integrating STEM Education and Humanities for Fostering Students’ Cultural Awareness Through CLIL Methodology. In: Anikina Z, editor. Integration of Engineering Education and the Humanities: Global Intercultural Perspectives. Cham: Springer International Publishing; 2022. pp. 405–414. doi:10.1007/978-3-031-11435-9_44

39. Gardner M. Beyond the Acronym: Preparing Preservice Teachers for Integrated STEM Education. AILACTE. 2017;14: 37–53.

40. Brownell SE, Price JV, Steinman L. Science Communication to the General Public: Why We Need to Teach Undergraduate and Graduate Students this Skill as Part of Their Formal Scientific Training. J Undergrad Neurosci Educ. 2013;12: E6–E10.

41. Illes J, Moser MA, McCormick JB, Racine E, Blakeslee S, Caplan A, et al. Neurotalk: improving the communication of neuroscience research. Nat Rev Neurosci. 2010;11: 61–69. doi:10.1038/nrn2773

42. Radford T. Of course scientists can communicate. Nature. 2011;469: 445–445. doi:10.1038/469445a

43. Cluley V. From “Learning disability to intellectual disability”-Perceptions of the increasing use of the term “intellectual disability” in learning disability policy, research and practice. Br J Learn Disabil. 2018;46: 24–32. doi:10.1111/bld.12209

44. Seewooruttun L, Scior K. Interventions aimed at increasing knowledge and improving attitudes towards people with intellectual disabilities among lay people. Res Dev Disabil. 2014;35: 3482–3495. doi:10.1016/j.ridd.2014.07.028

45. Kropp JJ, Wolfe BD. College Students’ Perceptions on Effects of Volunteering with Adults with Developmental Disabilities. Journal of Higher Education Outreach and Engagement. 2018;22: 93–118.

46. Kleinert HL, Jones MM, Sheppard-Jones K, Harp B, Harrison EM. Students with Intellectual Disabilities Going to College? Absolutely! TEACHING Exceptional Children. 2012;44: 26–35. doi:10.1177/004005991204400503

47. Skiba RJ, Simmons AB, Ritter S, Gibb AC, Rausch MK, Cuadrado J, et al. Achieving Equity in Special Education: History, Status, and Current Challenges. Exceptional Children. 2008;74: 264–288. doi:10.1177/001440290807400301

48. Connor DJ, Ferri BA. The conflict within: resistance to inclusion and other paradoxes in special education. Disability & Society. 2007;22: 63–77. doi:10.1080/09687590601056717

49. Elder TE, Figlio DN, Imberman SA, Persico CL. School Segregation and Racial Gaps in Special Education Identification. Journal of Labor Economics. 2021;39: S151–S197. doi:10.1086/711421

50. Rangvid BS. Special educational needs placement in lower secondary education: the impact of segregated vs. mainstream placement on post-16 outcomes. Education Economics. 2022;30: 399–425. doi:10.1080/09645292.2021.1995850

51. Lintangsari AP, Emaliana I, Fatmawati F, Rahajeng UW. Are students with disabilities ready for college? The influence of college readiness to college engagement. IJERE. 2021;10: 845. doi:10.11591/ijere.v10i3.21692

52. Rusch FR, editor. Beyond high school: preparing adolescents for tomorrow’s challenges. 2nd ed. Upper Saddle River, N.J: Pearson Merrill Prentice Hall; 2008.

53. Shogren KA, Ward MJ. Promoting and enhancing self-determination to improve the post-school outcomes of people with disabilities. Inge KJ, Wehman P, editors. JVR. 2018;48: 187– 196. doi:10.3233/JVR-180935

54. Beadle-Brown J, Murphy G, Wing L. Long-Term Outcome for People With Severe Intellectual Disabilities: Impact of Social Impairment. Am J Mental Retard. 2005;110: 1. doi:10.1352/0895-8017(2005)110<1:LOFPWS>2.0.CO;2

55. Skotko BG, Krell K, Haugen K, Torres A, Nieves A, Dhand A. Personal social networks of people with Down syndrome. American J of Med Genetics Pt A. 2022; ajmg.a.63059. doi:10.1002/ajmg.a.63059

56. Colleen TA, Lakin K, Carlson D, Domzal C, Austin K, Boyd K. Participation in Postsecondary Education for Students with Intellectual Disabilities: A Review of the Literature 2001-2010. The Journal of Postsecondary Education and Disability. 2011;24: 175–191.

57. Chris White, Nafilyan V. Coronavirus (COVID-19) related deaths by disability status, England and Wales: 2 March to 15 May 2020. Office for National Statistics. 2020.

58. Villani ER, Carfì A, Di Paola A, Palmieri L, Donfrancesco C, Lo Noce C, et al. Clinical characteristics of individuals with Down syndrome deceased with CoVID-19 in Italy—A case series. Am J Med Genet. 2020;182: 2964–2970. doi:10.1002/ajmg.a.61867

59. Villani ER, Vetrano DL, Damiano C, Paola AD, Ulgiati AM, Martin L, et al. Impact of COVID-19-Related Lockdown on Psychosocial, Cognitive, and Functional Well-Being in Adults With Down Syndrome. Front Psychiatry. 2020;11: 578686. doi:10.3389/fpsyt.2020.578686

60. Emes D, Hüls A, Baumer N, Dierssen M, Puri S, Russell L, et al. COVID-19 in Children with Down Syndrome: Data from the Trisomy 21 Research Society Survey. JCM. 2021;10: 5125. doi:10.3390/jcm10215125

61. Sabatello M, Landes SD, McDonald KE. People With Disabilities in COVID-19: Fixing Our Priorities. The American Journal of Bioethics. 2020;20: 187–190. doi:10.1080/15265161.2020.1779396

62. Landes SD, Turk MA, Formica MK, McDonald KE, Stevens JD. COVID-19 outcomes among people with intellectual and developmental disability living in residential group homes in New York State. Disability and Health Journal. 2020;13: 100969. doi:10.1016/j.dhjo.2020.100969

63. Rotarou ES, Sakellariou D, Kakoullis EJ, Warren N. Disabled people in the time of COVID-19: identifying needs, promoting inclusivity. J Glob Health. 2021;11: 03007. doi:10.7189/jogh.11.03007

64. Doody O, Keenan PM. The reported effects of the COVID-19 pandemic on people with intellectual disability and their carers: a scoping review. Annals of Medicine. 2021;53: 786–804. doi:10.1080/07853890.2021.1922743

65. Pellicano E, Brett S, den Houting J, Heyworth M, Magiati I, Steward R, et al. COVID-19, social isolation and the mental health of autistic people and their families: A qualitative study. Autism. 2022;26: 914–927. doi:10.1177/13623613211035936

66. Simmons AL. COVID-19 social distancing: A snippet view of the autistic social world. Disability & Society. 2020;35: 1007–1011. doi:10.1080/09687599.2020.1774866

67. Volkmar FR, Jackson SLJ, Hart L. Transition Issues and Challenges for Youth with Autism Spectrum Disorders. Pediatr Ann. 2017;46. doi:10.3928/19382359-20170519-03

68. Mazurek MO. Social media use among adults with autism spectrum disorders. Computers in Human Behavior. 2013;29: 1709–1714. doi:10.1016/j.chb.2013.02.004

69. van Schalkwyk GI, Marin CE, Ortiz M, Rolison M, Qayyum Z, McPartland JC, et al. Social Media Use, Friendship Quality, and the Moderating Role of Anxiety in Adolescents with Autism Spectrum Disorder. J Autism Dev Disord. 2017;47: 2805–2813. doi:10.1007/s10803-017-3201-6

70. Brugnaro BH, de Camargo OK, Corsi C, de Campos AC, Fernandes G, Pavão SL, et al. Functioning of children and adolescents with Down syndrome and the association with environmental barriers and facilitators during the COVID-19 pandemic. J Intellect Disabil. 2022;26: 824–838. doi:10.1177/17446295211032763

