## Supplemental Figures for "Experiential diversity training and science learning for college students alongside peers with intellectual and developmental disabilities"

| | |
|---|---|
| *"I want to learn more about adults with IDDs. I have only interacted with children with IDDs when I was younger, but I know I can* ***gain a new perspective*** *about disability from adults, I'm sure their life is completely different from mine."* | *"I want to be a teacher after college. I think [volunteering with LLWF] will foster creativity and the element of play when it comes to learning. I value* ***learning different teaching styles*** *and think this could provide a new perspective."* |
| *"I grew up a bit sheltered and unknowing of experiences beyond my own. I want to be able to* ***understand and empathize with other people's life experiences****. I also recognize how nice it is to have people to talk to and that adults with IDDs may not always have that opportunity. I want to provide that."* | *"Volunteering [with LLWF] will bring new friendships and new perspectives! I want to further my knowledge of the challenges adults with IDDs face. This will help me and others* ***foster a more inclusive and welcoming environment*** *in my future career."* |
| *"I volunteered at a rehab center and assisted at a physical and occupational therapy department for two years in high school. However, I almost never got to talk to the patients directly. I hope that volunteering will help me* ***learn to socialize and communicate with adults with disabilities****. I've never had this kind of experience in the past.* | *"I have had extremely limited experience with people with IDDs. I still aim to learn more about their lack of integration in society and in what ways I can* ***help advocate*** *for them. I want to get rid of any prejudice or bias I may unintentionally have towards people with IDDs."* |

**Supplemental Figure 1. Motivation for Volunteering with LLWF.** Representative quotes from pre-class survey. Prior to volunteering with LLWF, volunteers were asked about their motivations. Bold highlights text relating to benefits.

| | |
|---|---|
| *"I was really* ***surprised to see how social and interactive they were****. I thought that they might be shy in the beginning, but they were* ***open, friendly, and eager*** *to know about the volunteers and the class in general. They sometimes asked more questions than the UT students did!"* | *"I just didn't know anything about adults with IDDs. It was a question mark in my mind because I only remember having brief interactions with kids with IDDs when I was in K-12 because they weren't in our general classrooms. After getting to know the adults, I really enjoyed the reverse-inclusion format. They were* ***really welcoming and surprisingly enthusiastic*** *about participating in class."* |
| *"Before LLWF, I basically didn't know anything about adults with IDDs, so I wasn't exactly sure what to expect. I was pretty nervous coming in, but I've had such a great time being part of LLWF. Everyone I've gotten to know is so fun, sweet, and unique. The students all have such distinct personalities and I love hearing about what they have to say, whether it be about what they were going to do over the weekend to what they thought about the topics we learned about in class. They* ***share a lot of experiences that any other person would,*** *like romantic relationships, the struggle towards job searching, concerns about issues going around them, etc. I felt like I learned a lot when I talked with the other students, about them of course but also about myself."* | *"Meeting adults with IDD showed me* ***they're just as curious and enthusiastic about learning as adults without IDD****s and in some cases even more so because they don't really have the same opportunities as us. I didn't really understand what subjects' adults with IDDs enjoyed and didn't realize they could be* ***interested in diverse and sophisticated topics like everyone else****. I enjoyed talking with them about both lighthearted topics and heavy ones. In [one of the courses I volunteered for], we discussed some intense topics, such as crime, infidelity, and other mature content. My class peers with IDD were able to take in and discuss it just like the college volunteers could."* |
| *"[My expectations] changed a lot. I can see myself building genuine friendships with them. They are kind, thoughtful, and* ***try to achieve their fullest potential like everyone else."*** | *"I learned not to underestimate [people with IDDs] through volunteering. I realized* ***they want to talk and joke about things like I normally do with my friends****. I've learned that adults with IDDs are* ***just as capable as adults without IDDs****."* |
| *"I came to realize that* ***they are just like me****. A lot of us are the same age and have the same interests as me and my friends. I was surprised how everyone was welcoming and easy to talk to. I would've never known if I didn't volunteer. I didn't really understand what subjects' adults with IDDs enjoyed talking about. I didn't realize they could be as diverse as the general population—enjoying very lighthearted topics to very serious ones as well."* | **"***My expectations of adults with IDDs definitely increased after volunteering. The friends I made in class were smart, friendly, and insightful. People (including me, previously) may underestimate how self-aware [adults with IDDs] are. I learned that they are capable of critical thinking and intellectual growth just like any other student."* |

**Supplemental Figure 2: Changing volunteer's expectations of people with IDDs.** Representative quotes from volunteers in post-class surveys about how volunteering changed their expectations of people with IDDs. Bold highlights text relating to specific examples of changed expectations of people with IDDs.

| |
|---|
| *"I was one of the first volunteers in [LLWF over a decade ago]. I was fairly fresh out of high school, going to community college to get some credits out of the way before transferring to UT. **Volunteering in these classes helped to solidify the idea that I wanted to get my degree in special education**. Through [LLWF], I became connected to other disability organizations in Texas and began facilitating a community-based book club for adults with Down syndrome (I still do this today, ten years later). After graduating with my special education degree, I worked for 6 years teaching high school life skills in our local school district to students with disabilities. I now work at The SAFE alliance on their disability services team where my focus is providing healing classes to adults with disabilities and providing training to law enforcement on engaging with people / students with disabilities. **I see myself always being involved in some way in the disability community and it all really started with [Lifelong Learning with Friends]."*** |
| *"Through my time in the volunteering program, **I have become a more outgoing advocate for individuals with IDDs, an advocacy that has aided in my personal development and self-confidence**. [LLWF] has impacted the way I go about my research efforts, a topic that relates to ASD related genes and their behavioral implications. With the hands-on experience I am able to have with adults with IDDs, **I am able to contextualize the meaning of my research and the individuals I could potentially be improving the lives of.** Before volunteering with LLWF, I had wavering thoughts surrounding my career path. However, after being able to learn and connect with so many amazing adults from LLWF, many of whom demonstrated immeasurable kindness and new perspectives I would've never known otherwise, I know **I want to have a career that is centered around helping and improving the lives in the IDDs community**. The experiential learning that takes place goes beyond the academic topic of the class and lies within the experiences shared with my fellow classmates."* |
| *"My experience with the LLWF volunteer program has allowed me to learn about an exciting and fun new topic with students whom I would have never had the opportunity to collaborate with in my usual classroom setting. During small-group discussions, I have **learned how to be flexible with students, practicing patience and adaptability** with those who might be having difficulty with the topic at hand. Additionally, having conversations with my classmates has **benefited my social skills with peers who have unique needs**."* |
| *"I was shocked by how funny and fun they are to chat with. I found that they were all different and many could engage in deep and sincere discussions. More than that, they are great company. They're great listeners and are so genuine. This was such a **fun and easy way to volunteer on campus."*** |
| *"I think there are a lot of misconceptions about adults who live with intellectual and developmental disabilities. LLWF helped me to better understand a community of individuals who are as compassionate, if not more, about their education and autonomy [compared to neurotypical peers]. It's difficult not being able to go places on your own or have more barriers to communicating to the world. I've been fortunate to have [my experiences with] LLWF to be able to learn these lessons. This class also helped me understand how, even though there are resources in the community, it's not easy getting connected or staying connected to them. I hope that these lessons will help me in my medical practice in the future."* |
| *"I already had an awareness that adults with IDDs often have more limited opportunities, particularly when it comes to higher education, so I was immediately interested in learning more. I loved volunteering in these classes. I loved the opportunity to build relationships with adults with IDDs and other UT students. The reverse-inclusion model also really excited me; it was cool to be in these classes as a "volunteer" but also learning with the rest of the class. I'm not a 'science person', but [volunteering pushed me] to learn how to use different lab equipment, like microscopes and pipettes. "* |
| *"Volunteering with [LLWF] has been **my favorite part of undergrad.** To me, they have always been a rich source of conversation, friendship, and knowledge."* |
| *"I got connected to LLWF at the beginning of 2020, which was the first time I worked with adults with IDDs. **After I volunteered** for my first LLWF class, **I instantly knew this was a population I wanted to work with professionally**. I continued to volunteer for different LLWF classes and looked for more opportunities to work within the IDDs population. In early 2021, I started to work with a non-profit organization in Houston for adults with IDDs. I had the opportunity to shadow counseling sessions and hold check-ins with clients to gain more experience within the mental health field. After this experience, I knew this was the career path I wanted to take. Currently, I am a graduate student in clinical mental health counseling and plan to become a therapist for adults with IDDs. **Being a part of LLWF has opened a whole new career path for me** and has taught me the importance of continued support and education for adults with IDDs. "* |
| *"**LLWF was a valuable asset in broadening my professional career serving people with IDDs.** My plan was to go to medical school to specialize in neurodevelopmental disabilities, but my experience volunteering led me to explore the intersection of disability and education. Because of my positive experience with LLWF, I taught special education for a couple of years before entering medical school. That experience combined with my LLWF volunteering enriched my understanding of the like experience of people with IDDs. I'm sure this all will **help me relate better to my future patients."*** |

**Supplemental Figure 3. How LLWF Impacted Volunteer's Professional Aspirations.** Representative quotes from volunteers in post-class surveys from about how volunteering changed their professional interests. Bold highlights text relating to specific examples of LLWF impact.
